# Sustained multidimensionality of object representations as a scaffold for diverse behaviours

**DOI:** 10.64898/2026.09.17.752507

**Authors:** Amanda K Robinson, Sophia M Shatek, Tijl Grootswagers

## Abstract

The human brain transforms a brief glance into a neural representation flexible enough to serve many behavioural goals. This flexibility may reflect multidimensional visual representations read out selectively across tasks, rather than goal-specific instantiation. Combining a large-scale EEG dataset with a behavioural battery of 12 tasks spanning 200 objects, including perceptual, categorical and social decisions, we tested how behaviours are supported by stimulus-evoked representations. Information relevant to 11 of 12 tasks was accessible in the neural signal, emerging early and overlapping in time. Orthogonal behavioural dimensions showed distinct temporal profiles, indicating a genuinely multidimensional representation. The neural code converged on a human-relatedness axis, while feature-level information remained accessible. Behavioural accessibility was thus broad but structured. Object familiarity alone was reliable in behaviour yet absent from the shared neural signal, revealing a limit on what a glance makes accessible. We propose that sustained, recurrent processing serves this broad behavioural accessibility.

## Introduction

Visual perception feels effortless, yet a single glance at an object must support a wide range of perceptual and cognitive judgements. When we see an apple, we reliably identify it as an apple, judge its colour, shape and edibility, and evaluate possible actions towards it, even without explicit intent. This requires both stability and flexibility of visual perception: we must reliably recognise an object despite variations in lighting, viewpoint, size and context ^1^, yet the same object must support vastly different behavioural responses depending on our goals ^2^. Understanding how the brain constructs a representation that can support such diverse cognitive purposes is a central but unresolved question in visual neuroscience: how is information made available for whatever a situation demands, rather than only what is anticipated? To answer this, we need to characterise what a glimpse affords regardless of the task at hand; a floor on available information that any subsequent behaviour must build on.

Perceptual decisions span fundamentally different cognitive operations: some are fast and effortless, like detection, others slow and deliberate, like judging if something is ripe enough to eat; some rely on low-level visual features, others on meaning and affordance. Decisions can be made with only a glimpse of a stimulus ^3^, without prior planning of the task ^4^, and quick enough for only a feedforward sweep through the visual system ^5^. Flexible behaviour could be supported by neural object representations that are intrinsically rich and multidimensional, encoding diverse object properties rather than a single dominant dimension ^2,6–8^. Multidimensional representations could provide a substrate for diverse behaviours, without requiring a goal-specific representation for each. This multidimensionality may itself arise to serve diverse goal-driven behaviours ^9^, or could reflect the statistical structure of natural images ^10^, that may nevertheless influence downstream behaviours.

Multidimensionality is distributed rather than localised. Representational dimensions span both visual feature and categorical distinctions, with multiple interacting dimensions including animacy, real-world size, texture, shape, and colour ^8,11,12^. Although much of this work has focused on ventral visual cortex ^13–16^, multidimensionality also exists as early as primary visual cortex, which encodes position, retinal size, colour, and orientation for review, see ^11^. The distinction between low- and high-level processing is further blurred by the inherent correlation between visual features and object category ^17–19^. Indeed, visual features exhibit distinct but overlapping coding profiles across the prolonged temporal cascade of visual processing ^20^, suggesting that multidimensional visual representations are broadly distributed in space and time. Critically, this structure appears inherently behaviourally relevant, such that neural responses recorded during orthogonal tasks predict a range of perceptual judgements across diverse stimulus types, including visual similarity judgments for low-level stimuli ^20,21^, face authenticity judgments ^22^, and visual, conceptual, and face-likeness judgments ^23–28^. Further, large-scale similarity judgement work has revealed a rich high-dimensional space that relates behaviour and neural responses ^8,29,30^.

Because visual processing necessarily precedes the decisions it supports, the relevant information must be present by the time a decision is made. The open question is not whether the information exists, but when and how broadly it becomes accessible across processing. Prior work has established that object representations are inherently behaviourally relevant regardless of current task, but has largely concerned judgements that could be considered ecologically valid (e.g., categorising objects, judging similarity, detecting animals). The visual hierarchy, framed as progressively abstracting toward invariant high-level representations ^31,32^, is plausibly organised to support readout directly for such real-world tasks. Yet visual features can shape perceptual judgements ^19,33,34^, suggesting that early processing may be inadvertently used for behaviour too. To really test accessibility across the hierarchy, it requires judgements that target different levels of the cascade, from feature comparisons such as relative colourfulness to conceptual decisions about animacy. If accessibility tracks the hierarchy, correspondences should be temporally ordered, each judgement emerging as its information is computed. Alternatively, sustained recurrent processing may hold diverse information accessible across a prolonged window, so the same period serves all judgements regardless of which is required.

Sustained behavioural accessibility is plausible due to a separate, well-established observation: stimulus-evoked representations persist well beyond the feedforward sweep and even after the stimulus is no longer present. Objects are decodable for several hundreds of milliseconds, even under rapid, serial presentation and without any task requiring maintenance ^35–37^. This raises the question of why the visual system should hold information available by default. One possibility is that maintaining a broad representation enables a glimpse to serve whatever a situation later demands.

Here, we directly tested whether inherent object representations encode information relevant to diverse perceptual and cognitive judgements, and how that information is sustained over time. We combined a large-scale EEG dataset from previous work using 200 object stimuli ^36^ and existing behavioural dataset of three tasks ^25^ which we here enriched by creating a new openly available behavioural resource spanning nine additional tasks (Figure 1; Table 1). Together, these tasks cover rapid detection, perceptual feature judgments, categorical decisions, social judgments, familiarity, and unconstrained odd-one-out judgments under speeded and unconstrained conditions. Neural responses and behaviour were collected from different groups of participants as they viewed the same objects, yet performed separate tasks, ensuring that neural-behavioural correspondences reflect inherent stimulus-related rather than goal-directed representations.

**Figure 1.**
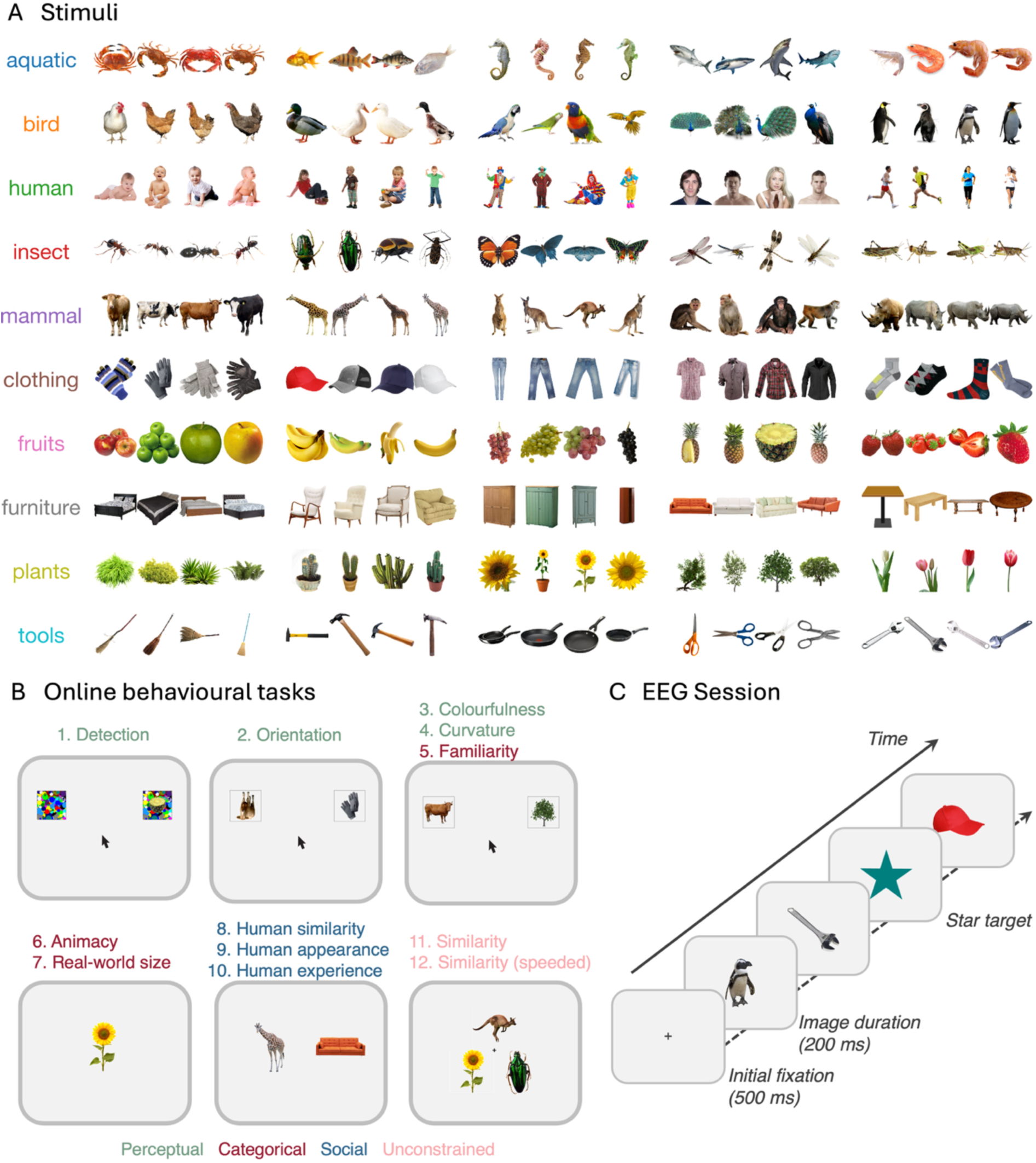
Experimental design. *Note.* **A)** Stimulus set consisting of 200 object images spanning 10 basic categories (5 animate, 5 inanimate), each with 5 object categories and 4 exemplars per category ^36^. **B)** Schematic of the 12 behavioural tasks used to probe responses to the 200 images. Tasks are numbered 1-12 and grouped as perceptual, categorical, social and unconstrained tasks. Each panel shows a representative trial display. The display is shown once where several tasks shared an identical display and differed only in the question asked. Full task instructions, stimulus presentation times, response measures and sample sizes are given in Table 1. **C)** Schematic of the EEG task ^36^, in which a separate group of participants viewed the same 200 objects in rapid serial visual presentation streams (200 ms per image, no gap) while performing an orthogonal target-detection task, ensuring neural responses reflect inherent rather than goal-directed object representations. Analysed data were from non-target trials.

**Table 1.** Overview of task characteristics.

|  | Task name | Task type | Instruction | Reference | Response | Measure | Stimulus presentation time | Subj Group | N | Reliability |
| --- | --- | --- | --- | --- | --- | --- | --- | --- | --- | --- |
| 1 | Detection | Perceptual | Which one contains an object? | Relative to a mask with no object | Click on a stimulus | Response time | Until response (Time out 1500 ms) | 1 | 107 | 0.664 |
| 2 | Orientation | Perceptual | Which one is more upright? | Relative (2 stimuli; 1 inverted) | Click on a stimulus | Response time | Until response (Time out 1500 ms) | 2 | 101 | 0.867 |
| 3 | Colourfulness | Perceptual | Which one is more colourful? | Relative (2 stimuli) | Click on a stimulus | Proportion chosen | Until response (Time out 1500 ms) | 2 | 101 | 0.984 |
| 4 | Curvature | Perceptual | Which one is more curved? | Relative (2 stimuli) | Click on a stimulus | Proportion chosen | Until response (Time out 1500 ms) | 1 | 107 | 0.965 |
| 5 | Familiarity | Categorical | Which one is more familiar? | Relative (2 stimuli) | Click on a stimulus | Proportion chosen | Until response (Time out 1500 ms) | 2 | 101 | 0.884 |
| 6 | Animacy | Categorical | Animate or inanimate? | Absolute | Key press | Proportion animate response | 200 ms | 3 | 86 | 0.923 |
| 7 | Real-world size | Categorical | Bigger or smaller than a shoebox? | Absolute | Key press | Proportion bigger response | 200 ms | 3 | 84 | 0.959 |
| 8 | Human similarity | Social | Which object is more similar to a human? | Relative (2 stimuli) | Key press | Proportion chosen | 200 ms | 4 | 44 | 0.835 |
| 9 | Human appearance | Social | Which object looks more similar to a human? | Relative (2 stimuli) | Key press | Proportion chosen | 200 ms | 5 | 38 | 0.870 |
| 10 | Human experience | Social | Which object thinks or feels more similar to a human? | Relative (2 stimuli) | Key press | Proportion chosen | 200 ms | 6 | 38 | 0.754 |
| 11 | Similarity | Unconstrained | Click the odd one out | Relative (3 stimuli) | Click on a stimulus | Pairwise proportion chosen | Until response | 7 | 96 | 0.930 |
| 12 | Similarity (speeded) | Unconstrained | Click the odd one out | Relative (3 stimuli) | Click on a stimulus | Pairwise proportion chosen | Until response (Time out 1500 ms) | 8 | 94 | 0.922 |

*Note.* Details for the 12-behaviour task battery. Tasks were grouped broadly as perceptual, categorical, social or unconstrained constructs. Participant group denotes which tasks were completed in the same session. *N* is final analysed sample size. Reliability is the mean Spearman-Brown split-half reliability, computed by splitting participants into two groups: For tasks 1-10, it is the split-half correlation of mean task performance (1,000 permutations), and for tasks 11 & 12, it is the split-half correlation between 1-dimensional scaling solutions of each dissimilarity matrix (10 permutations).

Many of these tasks required a relational or graded judgment about objects (e.g., which of two objects is more colourful, or whether an object is bigger than a shoebox) rather than discrimination of a low-level feature (e.g., is the object blue or red). Relational judgements integrate information across the object and compare it to another object or a stored reference, and so are not simply reducible to the presence of a single stimulus property of colour or size. Correlations between these judgments and the neural signal plausibly index whether the decision-relevant information is available in the representation, not just whether the underlying feature is encoded ^38,39^. We found that neural representations tracked behavioural responses across all but a familiarity task, with correlations emerging around 100 ms and sustained for up to 500 ms, and highly overlapping across tasks in time. We propose that sustained, recurrent processing may specifically serve behavioural accessibility, maintaining diverse task-relevant information independently of moment-to-moment demands.

## Results

### The task battery samples a rich behavioural space

The behavioural task battery encompassed 12 diverse tasks broadly separated into perceptual, categorical, social or unconstrained types, sampling judgements such as colourfulness, animacy, similarity to a human and visual similarity (Figure 1; Table 1). Participants performed the tasks as expected, for example animate stimuli elicited more ‘animate’ judgements than inanimate stimuli and humans were judged as more like a human than other stimuli on the humanness tasks. Yet, the pattern of behaviour across the 200 experimental stimuli varied markedly by task (Figure 2). For many of the tasks, behaviour separated largely by broad object category (e.g., in the similarity tasks, animacy, humanness tasks), but the category groupings varied by task. For instance, insects and mammals were grouped together on the animacy task, but separately on the real-world size task. Colourfulness judgements, by contrast, varied across stimuli within the same category. Tasks also revealed rich stimulus structure: upright orientation judgements were fastest for humans and mammals, possibly because these categories have canonical upright orientations. Fruit was judged most familiar, and furniture was judged least curvy of the broad categories. These differential judgements across tasks highlights that the same object can elicit a variety of judgements, and that behaviours can index distinct object dimensions rather than relying on a single categorical structure.

**Figure 2.**
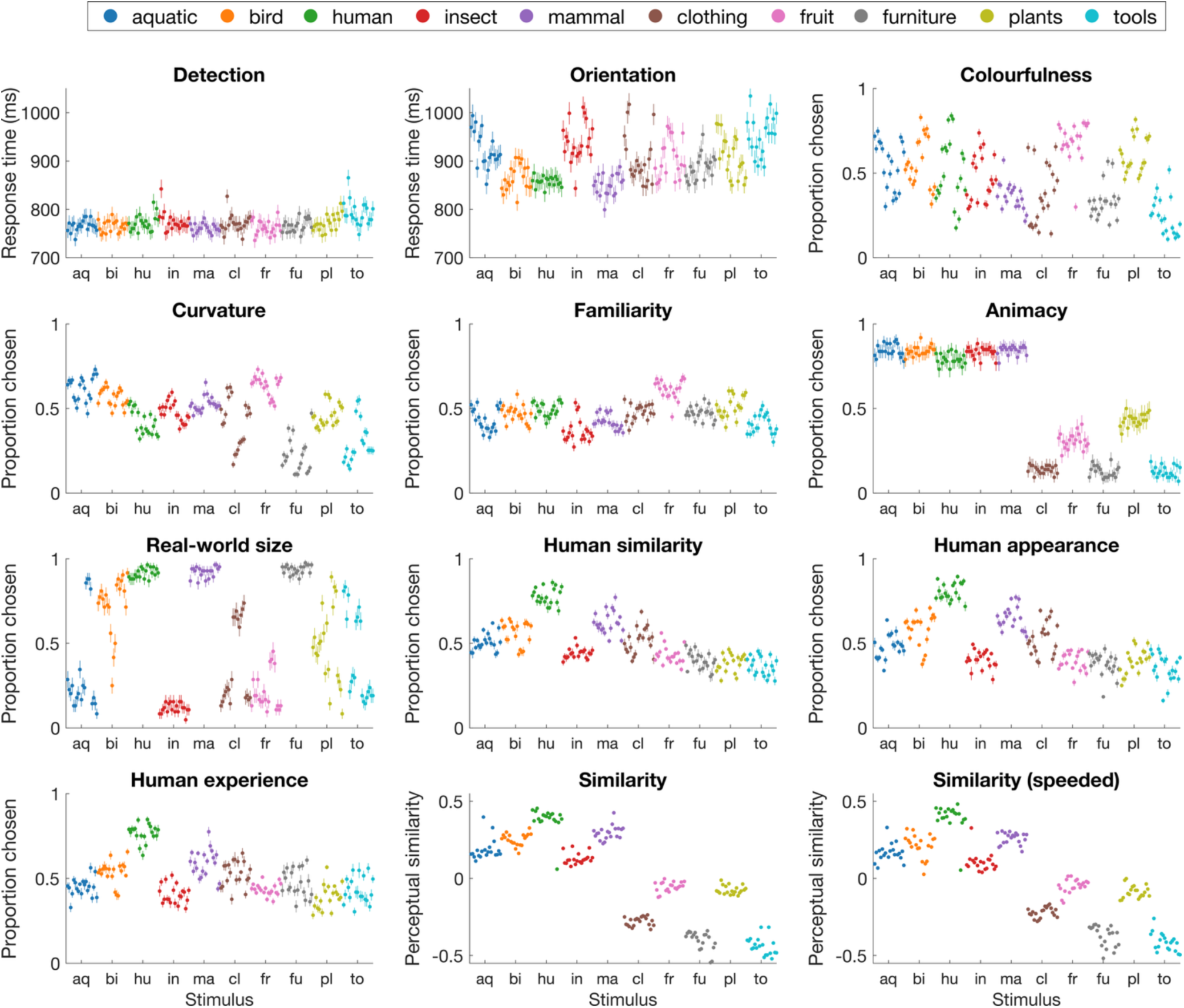
Behavioural results across 12 tasks show multidimensional responses to 200 stimuli. *Note.* Plots show relevant behavioural responses to 200 object stimuli (x-axis), colour-coded by object category. For detection and orientation tasks, which had ground truth correct answers, behaviour reflects mean reaction time. For the similarity tasks (untimed and speeded), stimuli are ordered by their position on a 1-dimensional scaling solution derived from odd-one-out choices, where stimuli positioned closer together elicited more similar behavioural judgements. For the remaining tasks (animacy, real-world size, curvature, colourfulness, familiarity and the three humanness judgements), behaviour reflects the proportion of trials on which each stimulus was selected as having the relevant property (e.g., more colourful, more familiar, more human-like). Error bars reflect standard error of the mean across participants.

### The behavioural space is multidimensional

To characterise the patterns of stimulus relationships across task, we computed a representational dissimilarity matrix (RDM) for each task by quantifying the pairwise similarity structure of behavioural responses across the 200 stimuli (Figure 3A). Tasks that evoked qualitatively similar patterns of behaviour, such as the two similarity tasks, showed similar RDM structure, while tasks tapping into different object properties (e.g., colourfulness versus familiarity) showed markedly different patterns. A two-dimensional multi-dimensional scaling (MDS) solution derived from the task RDMs reveals these relationships, placing tasks that evoke similar patterns close together in the task space (Figure 3B). The three humanness tasks seem to form a cluster, as do animacy, the two similarity tasks and curvature.

**Figure 3.**
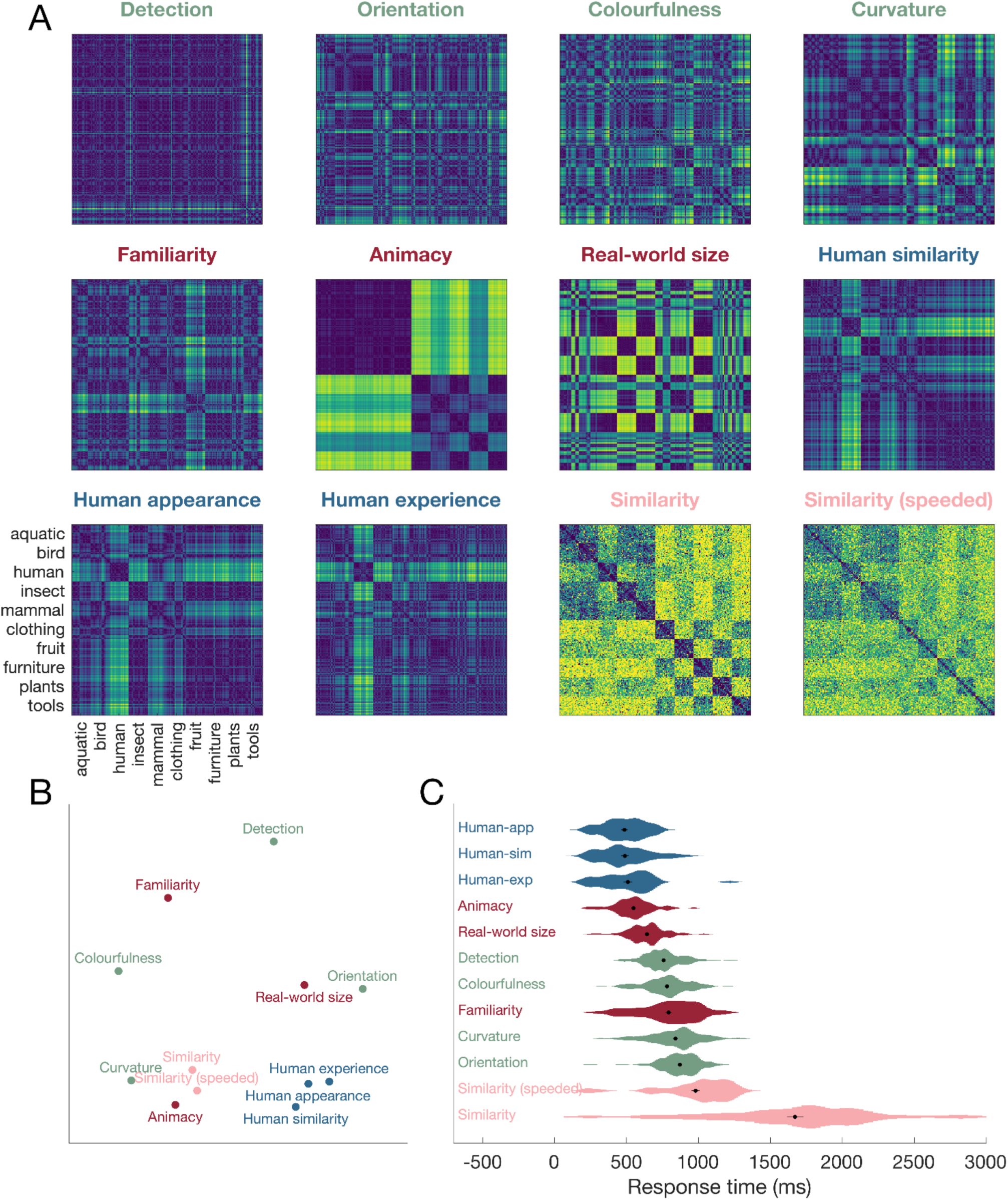
Relationship between different behavioural tasks. *Note.* A) Representational dissimilarity matrices (RDMs) for each task, showing the pairwise similarity structure of behavioural responses across the 200 stimuli. B) Two-dimensional MDS solution derived from the correlation between task RDMs, where distance reflects similarity in patterns of behavioural responses across stimuli; tasks closer together evoked more similar patterns of object similarity. C) Distributions of median reaction time per participant for each task. Black dots mark the mean across participants; error bars reflect one standard error. Tasks are colour coded by broad type: perceptual (detection, orientation, colourfulness, curvature), categorical (familiarity, animacy, real-world size), social (humanness tasks) and unconstrained (similarity tasks).

We also assessed response times per task. These varied substantially across tasks, ranging from approximately 500 ms for the humanness tasks, to 871 ms for the familiarity task, to over 1600 ms for perceptual similarity judgements (Figure 3C), likely reflecting differences in cognitive demands according to task question, reference frame (single image versus stimulus comparison) and response modality (mouse click versus key press). It should be noted, however, that tasks with similar paradigms and resulting RTs did not necessarily produce the same pattern of behaviour. Together, these patterns confirm that the 12 tasks sample qualitatively different aspects of object perception rather than redundant variations of the same judgment.

To quantify the dimensionality of the behavioural space and assess whether the 12 tasks captured genuinely independent dimensions of object variation rather than redundant information, we conducted a principal components analysis (PCA) on the 200 × 12 matrix of behavioural responses (Figure 4). Each column contained one task’s response profile across the 200 stimuli as in Figure 2, which was z-scored before running the PCA to equate tasks of different scales. If the tasks were largely redundant, for example if results on each task reflected the same categorical organisation of the stimuli, only one principal component would account for most of the variance; genuine multidimensionality would instead require several components to account for the variance across tasks.

**Figure 4.**
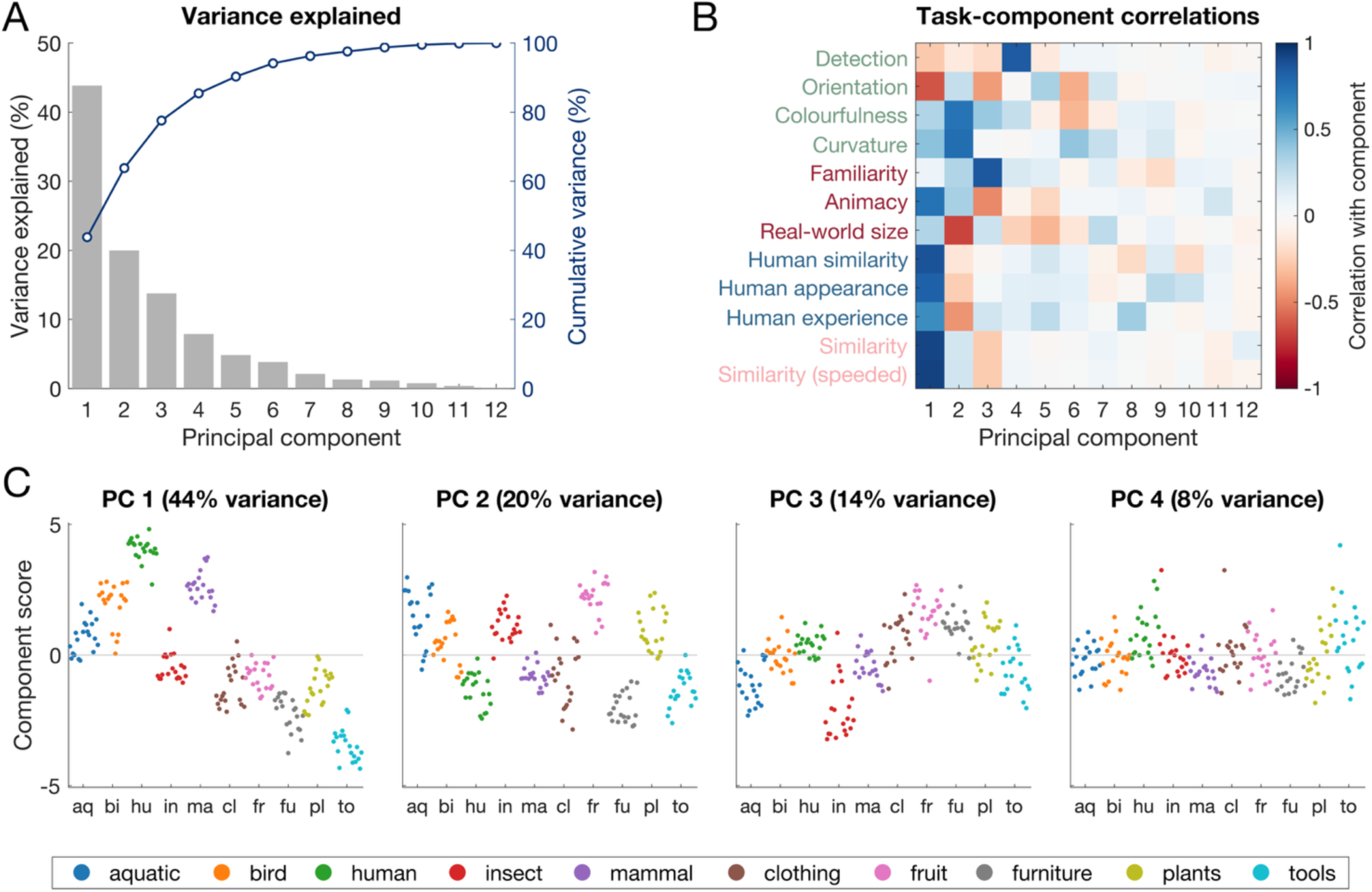
Principal components of behaviour across the 12 tasks. *Note*. PCA was conducted on a 200 (stimuli) × 12 (tasks) matrix of z-scored behavioural responses. **A)** Variance explained by each principal component (grey bars, left axis) and cumulative variance explained (navy line, right axis). The first seven components accounted for 95% of the variance. **B)** Spearman correlations between each task’s behavioural profile across the 200 stimuli and each PC’s scores. Each task was most strongly associated with one of the first four components. **C)** Scores of the 200 stimuli on the first four components, plotted in stimulus order and colour-coded by object category (legend below). Each component organises the stimulus set differently: PC1 separates stimuli along a broad human-similarity axis; PC2 generally separates small, rounded, colourful objects from large, angular, muted ones; PC3 is dominated by familiarity; and PC4, dominated by detection, shows little categorical organisation, likely varying with low-level image properties.

The PCA revealed the first seven components accounted for 95% of the variance (PC1: 44%, PC2: 20%, PC3: 14%, PC4: 8%, PC5: 5%, PC6: 4%, PC7: 2%; Figure 4A), indicating that the behavioural space is genuinely multidimensional but with tractable structure. Each of the 12 tasks was most strongly reflected in one of the first four components (Figure 4B). Projecting the 200 stimuli onto those components showed that each organised the stimulus set differently (Figure 4C).

PC1 (44%) captured a broad animacy axis, loading on the similarity tasks (ρ > .92), the humanness tasks (ρ = .62-.87) and animacy (ρ = .74), with orientation loading inversely (ρ = −.63; upright judgements were fastest for humans and mammals). The separation was not entirely binary along the animacy distinction; humans were most extreme, followed by mammals and birds, with insects falling among the inanimate objects, suggesting PC1 indexes human-similarity more than animacy per se. PC2 (20%) cut largely across this division, grouping aquatic animals, insects, fruit and plants apart from humans, furniture and tools. It loaded on curvature (ρ = .76), colourfulness (ρ = .73) and, inversely, real-world size (ρ = −.68), essentially separating small, rounded, colourful objects from large, angular, muted ones; stimuli varied markedly within category, indicating that this component tracks visual features rather than just object type.

The remaining two components were each dominated by a single task. PC3 (14%) was driven by familiarity (ρ = .86), with animacy loading inversely (ρ = −.48), grouping fruit, clothing and furniture, the categories regularly encountered in everyday life, separately from aquatic animals and insects. PC4 (8%) was associated almost exclusively with detection (ρ = .84; no other task exceeded |ρ| = .27) and showed less categorical organisation than the previous components. Objects that were similar on the categorical structure of PC1 were therefore often distant on the featural structure of PC2 or PC4, and vice versa. The 12 task behaviours thus sample a multidimensional space that cannot be reduced to a single principle. This provides the foundation for asking whether the neural cascade elicited by a brief glance simultaneously encodes information relevant to this full range of behavioural dimensions.

### Visual representations predict behaviour across diverse tasks

Having established that the 12 tasks tap genuinely different dimensions of behavioural variation, we asked whether time-resolved EEG responses during an independent target detection task predicted behavioural performance on each task using representational similarity analyses (RSA) ^7^. First, we constructed neural RDMs using decoding accuracy between all pairs of stimuli, repeated for each time point and participant. Then we correlated the neural dissimilarity matrices with the behavioural RDM for each task using Spearman correlation, and assessed statistical evidence of the group mean time-varying correlations using Bayes factors. Onsets were defined as the first three consecutive timepoints with BF > 10. To contextualise the magnitude of these correlations, we computed the lower noise ceiling, the between-participant reliability of the neural RDMs, which bounds the correlation any model could achieve ^40^. The ceiling peaked at ρ = .22 at 136 ms and remained above zero until ∼500 ms.

Neural representations were correlated with behavioural performance on 11 of the 12 tasks (Figure 5). Across tasks, correlations emerged within an early stage of processing. Onset time ranged from 92 ms (orientation) to 172 ms (human appearance), and most tasks showed reliable evidence between 100 ms and 200 ms (Table 2). This consistency is striking given the diversity of the tasks: the neural cascade encoded information relevant to perceptual feature judgements, categorical decisions, deliberate social judgements and unconstrained judgements at overlapping time periods, despite the variability in these responses and their reaction times.

**Figure 5.**
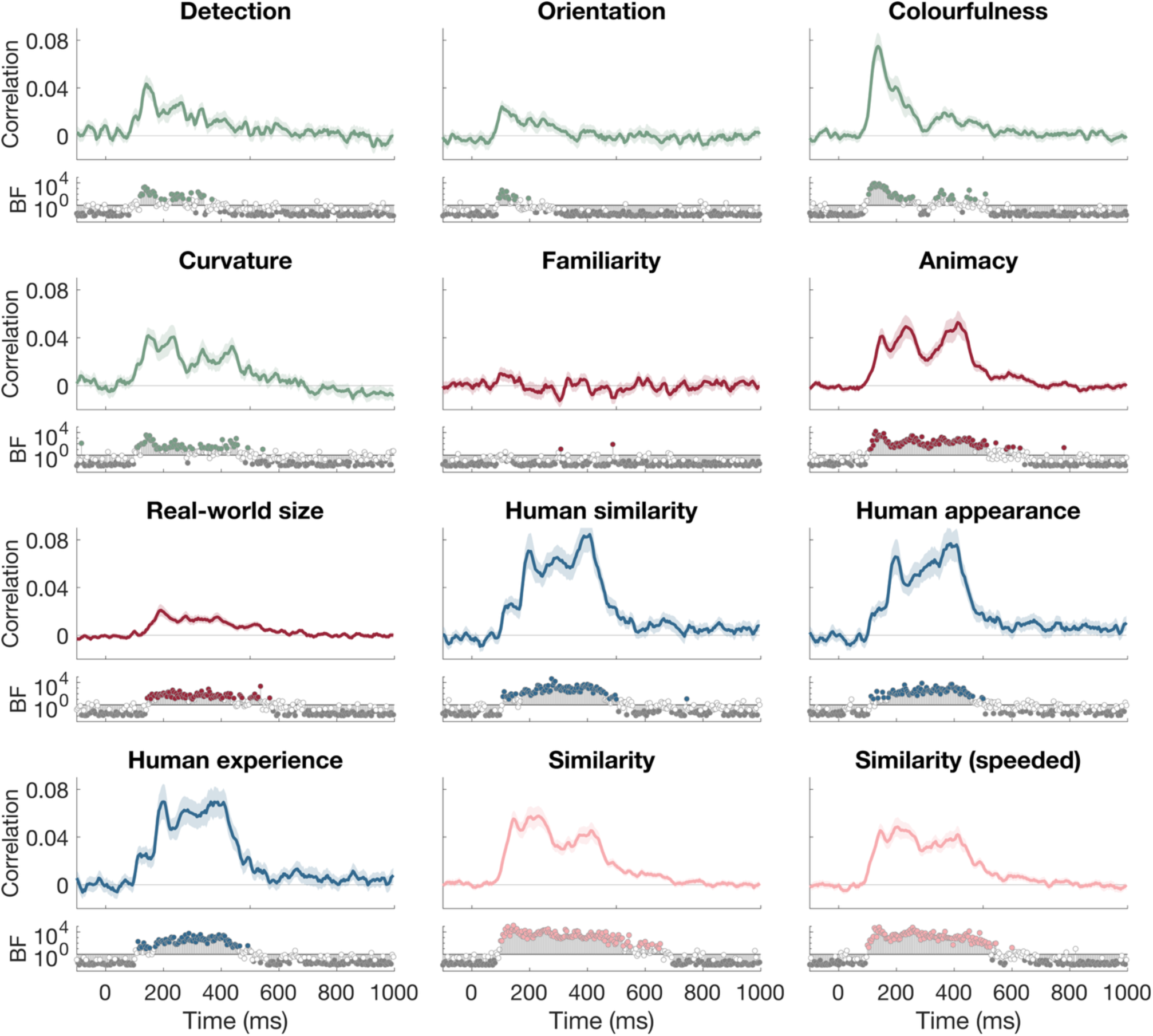
Neural time courses of relevant behavioural information across the 12 tasks. *Note.* For each task, the behavioural RDM was correlated with the EEG neural RDM at each timepoint, yielding a time course of the correspondence between neural representations and behaviour. Plots show mean Spearman correlations over time, and shaded regions indicate the standard error across participants. Below each time course, Bayes factors quantify evidence for correlations at each timepoint: filled coloured markers above the line indicate substantial evidence for an effect (BF > 10), open markers indicate insufficient evidence (10 > BF > 1/10) and grey markers indicate substantial evidence for the null (BF < 1/10). All tasks except familiarity showed reliable neural correspondence, with distinct onsets and peak latencies across tasks (see Table 2). Correlation traces are smoothed by 4 time points (16 ms) for visualisation.

**Table 2.** Characteristics of neural-behaviour correlations.

|  | Onset time<br>[95% CI] | #time points<br>BF>10 | Peak latency<br>[95% CI] | Peak<br>correlation | % of noise<br>ceiling at peak |
| --- | --- | --- | --- | --- | --- |
| Detection | 120 [88 132] | 35 | 136 [136 156] | 0.048 | 22 |
| Orientation | 92 [88 144] | 13 | 120 [96 244] | 0.026 | 14 |
| Colourfulness | 104 [92 108] | 48 | 136 [124 140] | 0.077 | 36 |
| Curvature | 108 [96 140] | 62 | 148 [140 432] | 0.043 | 24 |
| Familiarity | No reliable onset | 2 | 332 [12 904] | 0.013 | 17 |
| Animacy | 112 [100 124] | 107 | 416 [148 420] | 0.056 | 69 |
| Real-world size | 152 [92 216] | 80 | 184 [180 384] | 0.022 | 14 |
| Human similarity | 132 [100 184] | 88 | 412 [196 412] | 0.087 | 103 |
| Human appearance | 172 [100 208] | 81 | 380 [196 408] | 0.083 | 100 |
| Human experience | 112 [96 184] | 82 | 380 [196 412] | 0.076 | 92 |
| Similarity | 100 [84 108] | 125 | 224 [144 244] | 0.059 | 46 |
| Similarity (speeded) | 100 [96 112] | 110 | 196 [140 412] | 0.050 | 32 |

*Note.* Timing and strength of neural-behaviour correlations across the 200 stimuli for each task. All tasks except familiarity demonstrated reliable correlations with neural patterns of response. Onset time was calculated as the first of three consecutive time points with BF > 10. Confidence intervals were calculated by bootstrapping the group 1000 times with replacement.

Although absolute correlations were modest (peak ρ = .022-.087), they were substantial relative to the noise ceiling (max = .22). At their peak latencies, the humanness tasks reached the ceiling (>92 % of the reliability-adjusted maximum), indicating that the late neural representation was entirely organised by this social-similarity structure. Early-peaking perceptual tasks reached a smaller fraction of the high early ceiling (colourfulness 36%, curvature 24%, detection 22%), while real-world size and orientation were relatively low (both 14%). Yet, each effect was highly reliable, as indexed by the associated Bayes factors. Most tasks revealed overlapping and prolonged neural-behaviour correlations, and several tasks, notably the two similarity tasks and animacy, showed multiple distinct peaks, including an early one (∼200 ms) and a later one (∼400 ms), indicating that a single behaviour is supported by neural information at more than one processing stage.

Peak latencies, by contrast, were distributed across a broader time window, from 120 ms to 416 ms (Table 2). Critically, however, late onsets did not reflect late peaks. Information supporting animacy and humanness judgements was reliably present by 200 ms, yet was the strongest around 400 ms. Together, this suggests that rather than information relevant to different tasks appearing in sequence, they all appeared early and dynamically varied over time. Perceptual and feature-based tasks (orientation, detection, colourfulness, curvature) peaked relatively early (<150 ms), whereas tasks requiring categorical or social judgements (animacy, humanness) peaked later (real-world size 184 ms, others >380 ms). Despite this variation in peak timing, most correlations were sustained well beyond typical feedforward processing windows (>150 ms), and the time courses were broadly overlapping throughout.

### Familiarity is the exception

The familiarity task was the only task for which neural responses did not reliably predict behavioural performance. Bayes factors provided no sustained reliable evidence for a neural-behaviour correlation, with no reliable onset, and the peak correlation was negligible (ρ = 0.013 at 332 ms). This was not a matter of measurement reliability, because familiarity was itself a reliable, structured behavioural dimension, with notable stimulus organisation (Figure 3C), high split-half reliability (Spearman-Brown .884), and claiming its own principal component (PC3; Figure 4). This absence contrasts sharply with every other task tested and is therefore informative about the boundary of behavioural accessibility evoked by a stimulus.

### Behavioural dimensions are co-represented early, then collapse onto a categorical axis

Because the 12 tasks are correlated, the correspondence above partly reflects shared structure counted across multiple tasks. To separate the independent dimensions of behaviour, we correlated the neural signal with each orthogonal behavioural principal component. Applying the same onset criterion, eight of the twelve components were reliably tracked (top 4 PCs shown in Figure 6, all PCs in Figure S1). Most components had early onsets (PC1: 104 ms, PC2: 108 ms, PC4: 128 ms, PC6: 112 ms, PC7: 100 ms), and multiple orthogonal behavioural dimensions were co-represented from ∼100-170 ms. At the early stage the neural-behavioural correspondence was therefore high-dimensional: information relevant to many distinct behaviours was encoded concurrently, not in sequence. Confirming this, the total amount of variance explained in the neural data by the 12 principal components was higher than any individual component at the early time period (<300 ms; Figure 6). The variance explained by our behavioural battery as a percentage of the lower noise ceiling was 32% at 120 ms, 59% at 200 ms, 68% at 300 ms and increased to the maximum at around 400 ms, indicating a progression in the behavioural-relevance of visual representations over time.

**Figure 6.**
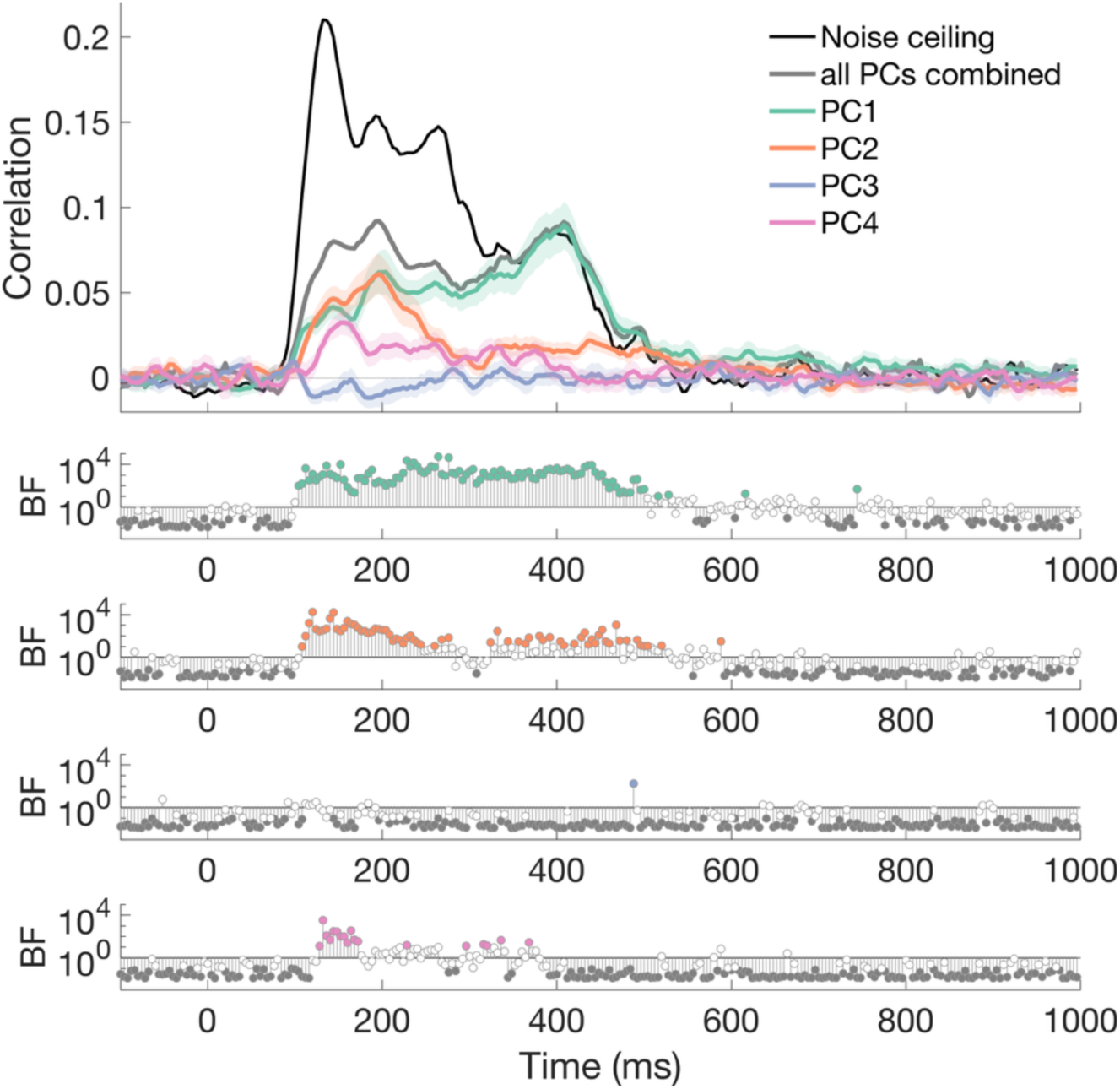
Correlation between neural representations and behavioural dimensions. *Note.* Plot shows the lower noise ceiling, calculated as the mean correlation between each participant with the rest of the group, compared to the total neural-PC relationship for all 12 PCA components of behaviour and each of the top 4 PCA components. PC1, PC2 and PC4 were all represented in an early time window, but PC1 solely accounted for the explainable variance from ∼350 ms. PC3, dominated by the familiarity task, did not reliably correlate with the neural geometry. Correlation traces are smoothed by 4 time points (16 ms) for visualisation.

These components differed markedly in persistence. PC1, the dominant categorical, human-similarity axis, was reliable for 104 timepoints, whereas no other component exceeded 68 (PC2) and most fell between 18 and 30. The transient dimensions decayed toward zero by ∼350 ms, whereas PC1 alone was sustained and strengthened, rising to a second, higher peak at ∼400 ms where it reached the noise ceiling. Over the epoch, the correspondence thus collapsed from a high-dimensional early representation onto a single dominant categorical axis.

Four components showed no reliable onset. Most notably, PC3, the familiarity-dominated dimension, was absent even in the early window, showing more support that familiarity’s behavioural dimension does not enter the neural signal at any stage. The other components (PC8, PC11 and PC12) explained variance in behaviour but were not reflected in the neural signal.

Categorical structure (PC1) was present from the earliest timepoints, simultaneous with perceptual and featural structure, rather than being derived from it later; and the early signal was not a single perceptual stage but a superposition of many behavioural dimensions at once. This suggests that what changes over the epoch is not which dimension is present but the *dimensionality* of the representation: an early, high-dimensional code in which information relevant to many behaviours is concurrently available, progressively resolving onto the categorical organisation that dominates late. Critically, because PC2 remained reliable well into the epoch before fading, perceptual and featural information stays available as categorical information accumulates. Thus, the representation elaborates rather than switches.

## Discussion

A single glance at an object is sufficient for a remarkable range of behaviours, from rapid detection to deliberate social judgements. Here, we asked whether the neural representation evoked by such a glance is relevant to behaviour across such diverse tasks. We show that a battery of twelve tasks performed on 200 object images sample a genuinely multidimensional behavioural space. Further, neural representations of the same stimuli during a different task tracked behaviour on eleven of these tasks, and the correspondences emerged early and were sustained for hundreds of milliseconds. The one exception, familiarity judgement, was itself a reliable behavioural dimension, revealing a boundary on what information is accessible from a glance. Together these results characterise behavioural accessibility as a broad but bounded property of visual object representation.

The central question we posed was whether behavioural accessibility is restricted to ecologically significant dimensions such as object identity and category, those that the visual system might be organised around ^1,32^. Alternatively, behavioural accessibility might extend to the full multidimensional structure of object representations, including feature-level information ^8,30^. Our results support the second, broader view. Neural representations predicted not only categorical and social judgements but also judgements of colourfulness, curvature, and detectability. A purely abstraction-driven hierarchy would be expected to discard these featural properties, yet we found they remained evident in the neural cascade. This extends evidence that explicit information about category-orthogonal properties increases rather than decreases along the ventral stream ^41^: such information is not only present in the representation, but accessible to behaviour.

Our behavioural task battery investigated a diverse range of perceptual, categorical, social and unconstrained tasks. Many of these tasks required graded, relational judgments rather than detection of a single feature. Judging which of two objects is most colourful, or whether an object is bigger than a shoebox, requires integrating information across an object and comparing it to another, or to stored knowledge. Such judgements are often assumed to be constructed at decision time, assembled by task-specific processes once the question has been posed. Our results suggest that much of what they depend on is already present before any question is asked. Colourfulness is a summary statistic computed over an entire object. Real-world size and human-relatedness depend on stored knowledge about objects rather than on image structure alone. Both types of judgement were nonetheless tracked by the evoked representation, early and under a task that asked for neither. Perceptual decision-making draws on several sources of information: prior knowledge about items, what the current stimulus provides, and the goal the observer is pursuing. A growing body of work emphasises the role of task in neural representations ^42,43^. Here, we characterised how perceptual decision-making operates on a baseline that is broad and shared across people. Thus, specific goals are plausibly used to reweight knowledge that is intrinsically evoked rather than creating it from scratch.

Diverse behaviours were supported by neural information with distinct temporal dynamics, but their correspondences emerged within a common early window and overlapped substantially in time, with most tasks reliably represented from 100-500 ms. This held even though the tasks differed in their response times by over a second. Behavioural diversity, even for the same objects, therefore appears to emerge not from task-specific neural states but flexible readout of sustained multidimensional representations. This offers a functional rationale for the persistence of visual representations far beyond stimulus duration ^35,36,44^: sustained, task-agnostic representations keep a rich set of object properties available so that they can be read out flexibly once a goal is specified, rather than requiring the system to anticipate every possible judgement or re-instantiate the stimulus after offset.

Decomposing behaviour into orthogonal dimensions revealed clear temporal structure. Early in the epoch, the neural signal reliably tracked many independent behavioural dimensions concurrently; a high-dimensional code in which information relevant to many behaviours was simultaneously available. Over the following few hundred milliseconds these dimensions diverged in persistence, with the perceptual and featural dimensions having relatively transient correspondences. The dominant human-relatedness dimension, however, was sustained and strengthened, rising to a late peak at which it accounted for all the reliable neural structure. Because these dimensions were rendered independent by construction (i.e., PCA yields orthogonal components), their divergent time courses indicate that different behaviours draw on distinguishable neural information rather than a single shared signal being relevant for every task. The behaviourally-accessible neural representation thus contracts over time from a high-dimensional early code onto a categorical organisation. Categorical dimensions are partly correlated with visual statistics, for example animals tend to be more curved than inanimate objects, which can make perceptual and categorical representations difficult to tease apart ^17,18^. Here, however, the perceptual dimensions peaked early (around 100 ms) and decayed, whereas the dominant categorical dimension peaked late (around 400 ms). A shared signal driven by common properties would not separate in time this way, so the two are unlikely to reflect the same underlying code. That the categorical dimension continued to strengthen after the image had been replaced in the RSVP task further suggests processing that outlasts a feedforward response. Both patterns are consistent with the sustained, recurrent processing we propose underlies behavioural accessibility, in which feedback and lateral computation maintain and elaborate an initial feedforward response while feature information stays available ^45–47^.

The contraction onto a human-relatedness axis connects our findings to a growing body of work identifying humanness as a prominent dimension of object representation ^25,48,49^. The separation we observed was not binary. Humans were most extreme, followed by mammals and birds, with insects falling among the inanimate objects. This fits recent arguments that the animacy organisation of occipitotemporal cortex reflects graded face and body likeness rather than a taxonomic distinction ^50–53^, and that animacy may be a misleading label for agency or humanness^54^. Our behavioural battery also lets us ask what distinguishes this dimension from the others expressed in behaviour. It is not the presence of a neural-behaviour relationship that is key, but its persistence. Referenced against the noise ceiling, individual behaviours captured a fraction of the reliable neural signal, with most tasks reaching 14-36%. Only the human-relatedness tasks approached the ceiling, and they did so late in time, when the neural structure was entirely organised by this categorical split. The same trajectory is evident when considering variance explained by the entire behavioural battery; together the behaviours explained around 40% early in time, but approached the noise ceiling later. Early on, then, the neural signal contains information that is not captured by the tasks here. This early residual structure may be due to information irrelevant to behaviour, or may reflect behaviours that we did not test.

This breadth of behavioural accessibility carries a deeper implication. Because so many diverse behaviours, including feature judgements that are tangential to canonical object recognition, could be predicted from the stimulus-evoked signal, the mere existence of a neural-behavioural correspondence is weak evidence that a dimension is central to how objects are organised in the brain. A significant correlation between behaviour and the neural signal demonstrates that the relevant information is accessible, not that the representation is organised around it. Concerns that findings in neural object representation may be overfit to particular stimuli or tasks ^17,49^ therefore also apply to these behavioural dimensions. Establishing that a dimension is genuinely central requires showing that the same scaffold generalises across stimulus sets and tasks.

The single exception to the generality of these effects shows the clearest boundary on behavioural accessibility. Familiarity was a reliable, structured behavioural dimension: it varied systematically across stimuli, claimed its own dimension in our decomposition, and was highly consistent across participants (split-half Spearman-Brown = 0.88). Yet, neural responses did not reliably predict familiarity judgements at any timepoint, and its corresponding dimension showed no reliable neural correspondence even early, where most other dimensions were tracked. Its absence is therefore not due to a noisy or unstructured behavioural measure. What distinguishes familiarity is the information it draws upon. A glimpse at an object certainly delivers what it looks like and what it is. These are the features that the other tasks draw upon: colourfulness, curvature and detectability relate to appearance, whereas animacy, human-relatedness and real-world size relate to meaning, drawing on stored knowledge. Familiarity is neither; it is the relationship between an object and experience with that object. It stands to reason that familiarity might modulate object representations, but itself isn’t a behaviourally-relevant feature evoked by the object. This is why familiarity can be highly agreed upon and still absent from the shared neural signal: people encounter similar things, so their judgements converge, but the property being judged is not one the visual system computes from the object. Thus, accessibility is bounded: what a glance makes available is general feature and categorical information that is evoked by the image, not how much experience a particular observer has had with the object.

Overall, here we provide evidence that a brief glance at an object generates a representation with information relevant to a wide range of behaviours. These types of information are available not in sequence but concurrently, early, and sustained for hundreds of milliseconds. Over processing, this representation sheds reliable multidimensional detail and converges on a human-relatedness organisation that comes to account for essentially all its reliable content, while earlier feature information remains accessible rather than being discarded. Yet this accessibility is bounded: what a glance makes available is general feature and categorical information about an object, not how familiar that object is to the individual. Behavioural accessibility, broad, sustained, and supported by recurrent processing, thus emerges as a fundamental but principled property of visual object representation.

## Method

A common stimulus set links all data in this study. EEG data collected with this stimulus set were drawn from previous work ^36^. We also include online behavioural data from three experiments in other prior work ^25^. For the current study, we collected new online behavioural judgements of the same stimuli across 9 different tasks. Raw and preprocessed neural data and stimuli are available online through openneuro: www.doi.org/10.18112/openneuro.ds004018.v2.0.0. Behavioural data and analysis scripts will be available upon publication.

### Participants

Across the EEG and 12 behavioural experiments, participants completed the tasks in return for course credit or payment. These studies were approved by the University of Sydney and Western Sydney University ethics committees. Informed consent was obtained from all participants.

We leveraged an existing EEG dataset from previous work ^36^, in which participants were 16 adults (5 females, 11 males; age range 18-38 years) recruited from the University of Sydney.

For the current study specifically, online behavioural judgements were collected for tasks related to perceptual similarity, object detection, object orientation, colourfulness, curviness, familiarity, animacy and real-world size. Separate groups of participants were recruited from the University of Sydney to complete different combinations of tasks: object detection and curviness (*N* = 110); colourfulness, orientation and familiarity (*N* = 113), perceptual similarity (*N* = 100); and time-limited perceptual similarity (*N* = 100). Another group of participants was recruited from Western Sydney University to complete animacy and real-world size categorisation (*N* = 101).

We also leverage an existing behavioural dataset from previous work ^25^. In this set, participants were recruited from Western Sydney University to complete three different online tasks relating to human-like judgements (*N* = 63; *N* = 65; *N* = 63).

Participants were excluded from analysis of a given behavioural task if they did not complete the majority of trials or their median response time was below 200 ms, indicating non-compliance with the task. After exclusions, the final sample sizes were: similarity (*N* = 96), speeded similarity (*N* = 94), animacy (*N* = 86), real-world size (*N* = 84), detection (*N* = 107), curvature (*N* = 107), colourfulness (*N* = 101), familiarity (*N* = 101), orientation (*N* = 101), like a human (*N* = 44), looks like a human (*N* = 38) and thinks like a human (*N* = 38).

### Stimuli

The stimulus set consisted of 200 visual objects from different categories ^36^. Briefly, stimuli comprised objects from two high level categories (animate, inanimate), within which there were 10 basic categories (5 animate, and 5 inanimate categories; e.g. mammal, tool, flower). Each of these 10 categories was further separated into 5 object categories (e.g., cow, dog, giraffe), and each object category consisted of 4 exemplar images (Fig. 1). This stimulus set comprises rich variability of visual features and conceptual categories, allowing considerable diversity in behavioural judgements.

### Behavioural experiments

Behavioural judgements were collected using online behavioural experiments ^55^ programmed in jsPsych ^56^ and hosted using pavlovia.org.

#### Perceptual similarity

Two experiments were designed to capture judgements of stimulus similarity. In each trial, three stimuli were displayed, and participants were asked to click on the odd one out (Figure 1B). Participants could use any criteria they wished to make their choice. A similar design has been used previously to model meaningful object geometry in large stimulus sets ^20,29^. In the first experiment, the images remained on the screen until participants made their choice. In the second experiment, participants had to make their choice within 1500 ms, after which the trial timed out. Each participant performed 400 trials each consisting of random combinations of stimuli.

To analyse the data, the similarity of each pair of stimuli was calculated (19,900 unique pairs for all combinations of the 200 stimuli). Experiment data were concatenated across participants. Pairwise similarity was computed as the number of trials where a stimulus pair was presented together but neither was chosen as the odd one out (i.e., how many times the third item was chosen as odd-one-out), as a proportion of the total number of trials that included the pair ^20^. Dissimilarity scores were calculated as 1 minus the similarity scores.

#### Object animacy and real-world size

Two experiments focused on the categorisation of the stimuli. For the animacy task, participants were presented with a stimulus for 200 ms and asked “animate or inanimate?”. They responded with the F or J buttons on the keyboard. In a separate block, participants completed the real-world size categorisation task. Images were presented in the same format as the animacy task but participants were asked “bigger or smaller than a shoebox?”. There were 400 trials in total, comprising one trial per stimulus and task combination. Task order and response keys were counterbalanced across participants.

Responses were analysed based on the mean response choice across the group per stimulus and task (i.e., proportion of ‘animate’ or ‘bigger’ responses per stimulus). Euclidean distance was calculated for each stimulus pair to result in a dissimilarity matrix per task.

#### Detection, orientation, colour, curviness and familiarity

These experiments were all two-alternative forced choice mouse click tasks. On each trial, participants first clicked a central “Next” button to initiate the trial. Two images (200 × 200 pixels) were then displayed, one to the left and one to the right of the screen, with their centres approximately 370 pixels either side of the screen midpoint and approximately 224 pixels above the starting cursor position. Participants clicked on the image that best satisfied the task criteria. Trials timed out after 1500 ms, followed by a 500 ms inter-trial interval. Breaks were offered every 50 trials.

In the detection experiment, we assessed object detection judgements using a two-alternative forced choice task. Two Mondrian masks ^57^ were presented, and one was overlaid with an experimental stimulus (Figure 1B). Participants were asked to click on the item that contained an object. There were 400 trials, two for each of the 200 stimuli (one presented per left/right visual field).

The orientation experiment was designed to assess judgements of whether an object was upright or inverted. Participants were presented with two objects, one upright and one rotated 180 degrees with respect to the original orientation in the stimulus set (Figure 1B). They were asked “which one is more upright?” and had to use the mouse to click on their choice. There were 400 trials across the experiment, with each object being presented twice in an upright and twice in an inverted orientation. Stimuli were presented in random combinations, with the exception that the upright and inverted versions of the same stimulus could not be presented together.

For colour, curvature and familiarity, in each trial, participants were presented with two stimuli and had to click on one that best satisfied the criteria (Figure 1B). They were asked “which one is more colourful?”, “which one is more curvy?”, and “which one is more familiar?”. There were 400 trials for each task, with four repeats for each of the 200 stimuli (two presented per left/right visual field).

For detection and orientation, when there was a ground truth correct answer, responses were analysed based on median reaction times for each stimulus per participant, and the mean taken across the group. Only upright presentations were analysed for the orientation task.

For colour, curvature and familiarity, responses were analysed based on mean response per stimulus per participant (i.e., proportion of time the stimulus was chosen relative to the number of times it was shown), then the group mean response was calculated for each stimulus.

Dissimilarity scores were calculated as the Euclidean distance between stimulus estimates (mean reaction time or mean response) for each pair of stimuli.

#### Humanness

Data from three experiments were drawn from previous work ^25^ to assess the perceived similarity of stimuli to humans. On each trial, a fixation cross was presented for 500 ms, followed by two stimuli for 200 ms. The two stimuli (224 × 224 pixels) were presented side by side with a central fixation cross between them, centred on the screen. Participants were asked either “which object is more similar to a human?”, “which object looks more similar to a human?”, or “which object thinks or feels more similar to a human?”. They responded using the F and J keys on the keyboard to correspond with a left or right image choice, respectively. Each experiment used a single randomly selected exemplar from each of the 50 object categories, resulting in 1225 trials per participant. After all participants were combined, this covered all 200 stimuli. Breaks were offered every 100 trials.

Responses were analysed based on the proportion of response choices across the group per stimulus and task. Euclidean distance was calculated for each stimulus pair to result in a dissimilarity matrix per task.

### Multidimensionality in behavioural judgements

To assess the common dimensions in results across the 12 tasks, we conducted principal components analyses (PCA). For each task, we used a 200 x 1 vector of behaviour across the 200 stimuli. For detection and orientation, the measure was response time. For colour, curviness, familiarity, animacy, real-world size, and the three humanness tasks, the measure was mean response choice. For similarity and speeded similarity, we converted the RDMs into 1-dimensional vectors using multi-dimensional scaling. Each of the 12 task vectors was z-scored and then all vectors were concatenated before running PCA. This resulted in 12 principal components (PCs), each accounting for different amounts of variance in the behavioural data. To assess the information within each PC, we calculated Spearman correlation between each PC and the original data per task.

### EEG data and analysis

EEG data are described in Grootswagers, Robinson, and Carlson (2019). Electroencephalography was measured while participants viewed all 200 object stimuli in rapid serial visual presentation sequences. Here we focus on the data when images were presented at 5 Hz (200 ms per image, no gap), which was the longest presentation time that resulted in the strongest stimulus representations in the original study. Stimuli were presented in random order with 1-4 embedded targets (boats or stars) as an attention task. Participants counted targets and responded at the end of each sequence. Crucially, the 200 images analysed here were non-targets in the experiment.

To assess the dynamic neural geometry associated with the 200 objects, we used time-resolved neural decoding of images, collapsed across the two tasks, as in the original paper ^36^. EEG data were filtered with a high-pass of 0.1 Hz and low-pass of 100 Hz, and epoched between −200 to 1000 ms relative to every image presentation. Neural decoding was conducted for every pair of objects (i.e., 19,900 pairs for the 200 objects). For every time point, we used leave-one-sequence out cross-validation with a linear discriminant analysis (LDA) classifier to obtain time-varying decoding accuracy for each pair of images. This decoding accuracy indexed the visual information contained in the EEG signal across all object images, allowing us to assess the geometry of neural representations and relate it to different types of behaviour.

### Analysis

We used Representational Similarity Analysis (RSA) ^14^ to assess how the behavioural patterns related to the neural representations to the same 200 stimuli. For the neural data and each of the behavioural tasks, we modelled representational dissimilarity for the 200 objects, which abstracted away from the actual measure (e.g., neural decoding, reaction time, accuracy, behavioural principal components) and allowed us to correlate between different measures. Mean RDMs were collated for each behavioural task across participants. Neural RDMs were collated for each time point separately for each participant. Time-varying neural-behaviour relationships were assessed by using Spearman correlation to compare the group behavioural RDMs with neural RDMs of each EEG participant per time point, then calculating the mean across the group.

This same neural-behaviour correlation method was used to relate each of the 12 principal components based on behaviour, modelled as an RDM using Euclidean Distance, to the neural representations. To quantify the total variance in the neural representations explained by the combination of behavioural components, we used the same 12 principal component RDMs as joint predictors of the neural representations. At each time point, the predictor RDMs were entered into a linear model and fit to the neural RDM using leave-one-participant-out cross-validation, with ordinary least squares weights estimated from the mean neural RDM of the training participants, used these weights to generate a predicted neural RDM, and then correlated (Spearman) this prediction against the RDM of the held-out participant. The cross-validated correlation was averaged across participants to give the total explained representational similarity at each time point.

As an estimate of the highest correlation attainable given the variance in the neural data, we estimated the lower bound of the noise ceiling ^40^ at each time point by correlating (Spearman) each participant’s neural RDM with the mean neural RDM of the remaining participants, again using a leave-one-participant-out procedure, and averaging across participants.

For each behavioural task, we characterised the timing of the neural–behaviour relationship by estimating the onset and peak of the correlation time course. Statistical evidence at each time point was quantified with Bayes factors (see below). The group onset was defined as the first time point at which the Bayes factor exceeded 10 for three consecutive time points, and the group peak as the time of maximum mean correlation. To obtain confidence intervals on these estimates, we bootstrapped across participants (1000 resamples with replacement), recomputing the Bayes factors, onset, and peak on each resample; onsets were searched from 50 ms onwards as anything earlier than this is implausible due to delays in signal transduction. 95% confidence intervals were calculated from the bootstrap distributions.

### Statistical inference

We used Bayesian hypothesis testing to quantify the evidence for the alternative relative to the null hypotheses ^58–62^. Bayes Factors (BFs) quantify evidence continuously, so do not call for explicit correction for multiple comparisons. Bayes Factors were calculated using the ‘BayesFactor’ package in R ^63^ using a JZS prior on effect size with default scale r = .707 ^60,61,64,65^. The null hypothesis was specified as an interval of standardised effect sizes between −0.5 to 0.5 centred on a null value of zero ^66^. A Bayes Factor expresses the marginal likelihood of the data under the alternative hypothesis relative to the null hypothesis. Following common conventions, we treated BF > 10 as strong evidence for the alternative hypothesis, and BF < 1/10 as strong evidence in favour of the null ^60,67^.

