## Supplementary figures and images for "Sustained multidimensionality of object representations as a scaffold for diverse behaviours"

### Supplementary Figure

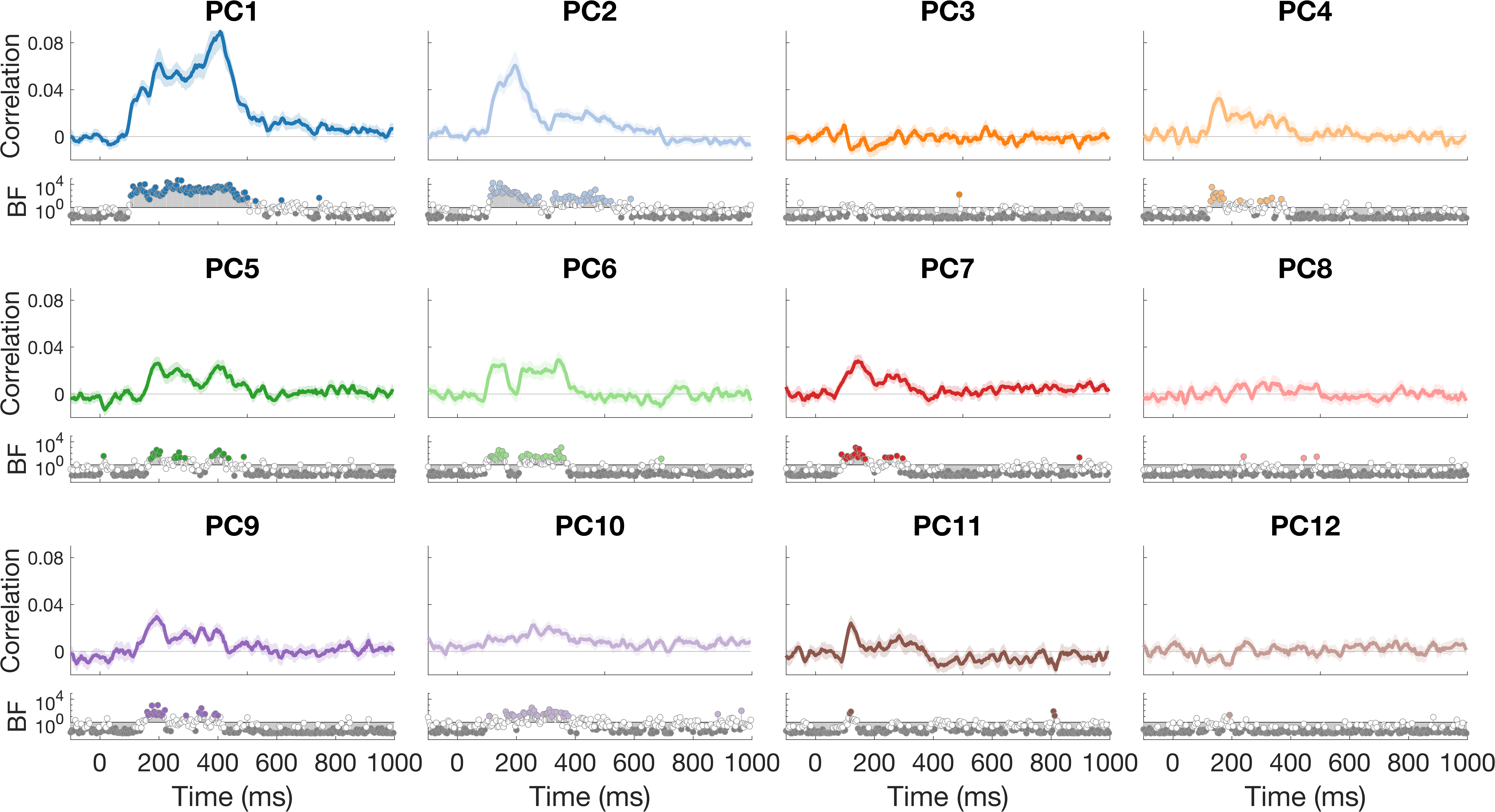
